# Digitoids: a novel computational platform for mimicking oxygen-dependent firing dynamics in *in vitro* neuronal networks

**DOI:** 10.1101/2023.10.30.564673

**Authors:** Rachele Fabbri, Arti Ahluwalia, Chiara Magliaro

## Abstract

*In vitro* models of neural tissues are crucial to gain new insights on the pathophysiology of the brain. However, cell cultures are associated with many drawbacks and difficulties, e.g., technical complexities, ethical problems and high cost. Computational model-based solutions could represent an important tool to support the study of neuron function. In this work we present a novel computational platform where digital neuronal networks, i.e., Digitoids, can be developed with different size and layouts. The Digitoids rely on a novel firing model where the dependence on oxygen concentration is introduced, since it is a crucial limiting factor in cell cultures. To validate the performance of the platform, Digitoids were developed with the same morphological arrangement as observed in neuron monolayers *in vitro*. The comparison between the functional output of the Digitoids and the experimental data are not statistically different. The platform delivers a flexible digital tool that can easily be adapted for mimicking *in vitro* models with increasing complexity and can be exploited to optimize the laboratory experiments involving neuron cultures.

**Author Summary:** The use of cell models within laboratories is crucial to gain new understandings of the functioning of neuronal assemblies. However, culturing cells requires highly skilled personnel, a huge quantity of disposable materials and is thus associated with elevated costs. To overcome these limitations, the use of computer-based systems that reproduce the electrical behaviour of neurons can be employed. Since *in vitro* models are vessel-free, we present a novel computational model of neuron electrophysiology, where we introduce the dependence on local oxygen concentration Indeed, oxygen affects the metabolism and the function of cultured neurons. Thanks to our model and platform, we are able to reproduce the morphology of the cultured networks of neurons and build the so-called *Digitoids*. We then simulate the *Digitoids* and obtain their electrophysiological output. We compared the *Digitoids’* output with experimental data from neurons cultured on microelectrode arrays. We did not find any differences between the electrophysiological output of the *Digitoids* and the experimental data, therefore we conclude that our novel oxygen-dependent model can be used to develop more physiologically relevant tools for simulating the activity of cultured neurons.

## 1. Introduction

*In vitro* models of neural tissues are emerging as a promising, more ethical and accessible way to study brain pathophysiology with respect to animal models. Indeed, their high controllability and observability have allowed the investigation of neurotoxicity, neuroprotection, drug screening and therapeutic assessment for different neuropathies [1–3].

The electrophysiological behaviour of cultured neuronal networks can be studied via patch-clamp, calcium imaging and microelectrode arrays (MEAs) [4]. None of these technique allow both a minimally invasive and a complete characterization of the electrophysiological activity. Indeed, each method comes with its key features and advantages but also drawbacks and limitations in terms of analysis resolution (both spatial and temporal), invasiveness and technical complexity, as summarized in Table 1. In patch-clamp techniques, cell activity is closely monitored with single-channel event resolution, also capturing the subthreshold membrane voltages [5], but it is a complex and extremely invasive technique, allowing the analysis of a restricted number of neurons and only for a limited timespan [6]. On the other hand, calcium imaging employs fluorescent protein sensors to record intracellular calcium activity, which is an indirect indicator of neural activity [4], but for technical limitations in microscopy, it is not capable of monitoring a neural population with the large spatial extent of the MEAs and has relatively poor temporal resolution [7]. For the MEA technology, neurons are usually cultured on the electrodes, which collect extracellular field potentials generated from the activity of the closest neurons [6]. Thus, MEAs do not affect cells and can be executed long-term, up to several days and months [8]. As such, they are the most widely used and effective methodology for monitoring *in vitro* network electrophysiology at high temporal resolution [6,8]. Despite that, for technical drawbacks (mainly due to electrode size, their distributions and ability to collect the electrical signals from the neurons nearby), MEAs recordings cannot provide a precise characterization of single-cell behavior [9].

**Table 1.** Advantages and disadvantages of the patch-clamp, calcium imaging and microelectrode arrays techniques.

| Technique | Advantages | Limitations | Refs |
| --- | --- | --- | --- |
| **Patch-clamp** | Single cell recording; single channel resolution; subthreshold activity monitoring | Complex technique requiring highly skilled personnel; invasive and cell-damaging (not allowing repeatable experiments); analysis limited to few neurons; limited timespan analysis | [4,6,10,11] |
| **Calcium imaging** | Single cell and small groups of cells recording; synaptic transmission and plasticity mechanisms analysis | Unable to capture complex dynamics of the whole-network; limited timespan analysis | [4,12] |
| **Microelectrode arrays** | Non-invasive; simultaneous whole-network recording (with spatial resolution depending on the device); very long timespan analysis (up to several months); repeatable experiments allowed; relatively easy to use | Unable to provide single cellular and subcellular information; sparse recording of extracellular field potentials; handling and processing the high amount of raw data produced | [4,6,8,13,14] |

Neuronal cultures are a low-throughput technology, which needs highly skilled personnel, and the reagents used for their generation and maintenance are expensive [15,16]. Moreover, they need a sterile environment, incubators, and high volumes of single-use plastics, all having a negative impact on climate and sustainability [17]. Over the past years some computational model-based solutions for generating virtual representations of neuronal cell cultures which replicate most of the experimentally observed salient firing properties have been proposed [18]. Intuitive and easy to use simulators (e.g., BRIAN 2, NEST, NEURON) have been employed to efficiently simulate spiking neural network models [19–21]. Although they have not been purposely implemented for mimicking *in vitro* dynamics, they have been successfully exploited, e.g., to study neuronal network modulation and delineate potential mechanisms underlying activity patterns [22]. Traditionally, these models include the mathematical descriptions of the single-neuron activity, ranging from simple phenomenological characterization of neuronal spiking [23] to more complex, conductance-based simulations of ion fluxes between the intra and the extracellular space [24], as well as of the cell-cell connections to resemble the neuronal network architecture [25]. Some studies also incorporate more sophisticated models, e.g., including astrocytes via tripartite synapses [26].

However, few of these approaches are based on energetic considerations, i.e. the dynamics of oxygen consumption or ATP hydrolysis. Beyond the basic activities common to other cells (e.g., DNA and RNA synthesis), resource uptake in neurons is also dedicated to support spiking activity, because of the role of the Na^+^-K^+^-ATP pump in signal propagation [27,28]. Since ATP dephosphorylation depends on the rate of oxygen consumption, oxygen dynamics can be monitored [29,30]. Indeed, oxygen is crucial in *in vitro* cultures, and particularly so in 3D cell-laden models (e.g., neurospheres, organoids), where the lack of an adequate network for nutrient supply results in concentration gradients within the constructs [31,32]. On the other hand, in traditional monolayers cells are inevitably exposed to different oxygen concentrations for the different culturing conditions, i.e., the amount of medium or the oxygen boundary concentrations [33–36]. An analytical formulation describing oxygen-dependent firing in tissue slices was proposed by Wei and collaborators in 2014 [37] to elucidate the mechanisms of seizure development and termination as well as the interaction between seizures and energy metabolism. Their model assumes that oxygen variations around the neuron depend on the diffusion from the bath solution and on the consumption rate for firing, but do not consider that oxygen can also be consumed for sustaining other metabolic functions of the cell.

Starting from Wei et al.’s model, we have developed a computational platform able to mimic *in vitro* electrophysiological behaviour at the single-neuron and network level. Such digitalized versions of *in vitro* neuronal monolayers, where the dependence on local oxygen concentration is considered, are called *Digitoids*. As *in vitro* networks can have very different culture conditions and layouts in terms of number of neurons, their densities and their connections [38–41], the platform is purposely designed to be modular, thus the user can generate *Digitoids* matching any type of *in vitro* neuronal network [36–39]. For testing the performance of the *Digitoids* and highlighting the crucial role of oxygen in accurately describing neuron dynamics observed *in vitro*, we digitalized the layouts of neuron monolayers seeded on commercial MEAs. Then, the graphs describing the *in vitro* layouts were used to build the corresponding *Digitoids* with an oxygen-dependent model of firing and metabolism. Additionally, traditional network models neglecting the oxygen dependence on firing activity were implemented as a control to compare against the *in vitro* recordings and assess the degree of similarity between the *Digitoids*’ outputs and the experimental data.

## 2. Outline of the computational platform

The computational platform was developed in Matlab (the Matworks Inc., Boston Massachusetts), exploiting the Simulink toolbox. In the following sections, the geometry of the *in vitro* monolayer, how it can be replicated within the platform to obtain the model of the single neuron dynamics and the creation of the neuronal networks, i.e., the *Digitoids*, are described.

### 2.1 The Geometry

Fig 1.a shows the geometry of the system to be modelled, composed of a well, seeded with cells homogenously distributed on the bottom and filled with a layer of culture medium of a specific height *h*. The neurons form synaptic connections to create the neuronal network [42,43]. The top layer of the medium at the interface with air maintains a uniform and time-invariant oxygen concentration *c*_0_. Its value is typically 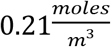but can be set to a defined value according to the experimental setup to be reproduced.

**Fig 1.**
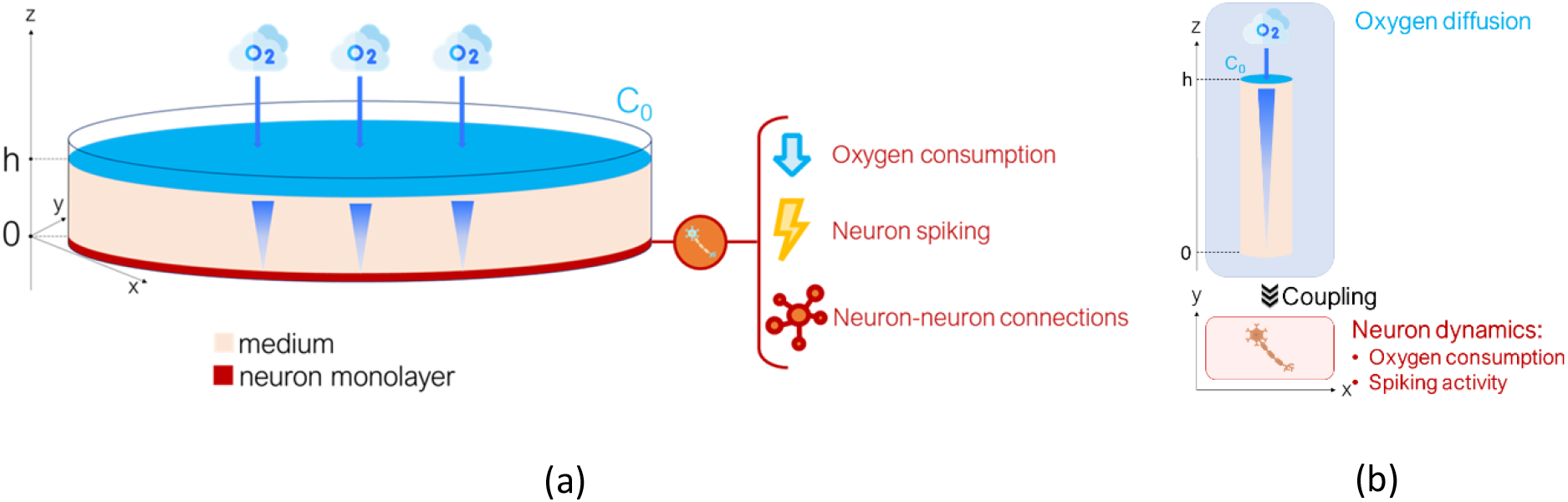
Geometry of the modelled system. a) The in vitro system with all the phenomena involved: oxygen diffusion (from the top level of the medium z=h to the neuron z=0), oxygen consumption by the neuron, neuron spiking and network dynamics (i.e. neuron-neuron connections); b) Sketch of the simplified system modelled in the platform.

In such a system, four phenomena are involved: i) oxygen diffusion through the medium, ii) oxygen consumption by neurons at the bottom of the well which generates the metabolic energy for fuelling cell vitality and function, iii) single-neuron spiking activity and iv) neural network dynamics, i.e., the transfer of electrical information via cell-cell connections.

Given the characteristic times of the four phenomena involved, reported in Table 2, we can assume that the passive diffusion of oxygen is independent of the x and y directions and only occurs along the z axis [47–49]. Moreover, while the electrophysiological activity of the single neuron is oxygen-dependent, the dynamics of the network is mainly influenced by the strength of neuron physical connections. Given these considerations, the whole neuronal network (Fig 1.a) can be described as the interconnection of single neuron entities (as the one depicted in Fig 1.b), where oxygen diffuses along the z axis and reaches the cell, that consumes it.

**Table 2.**
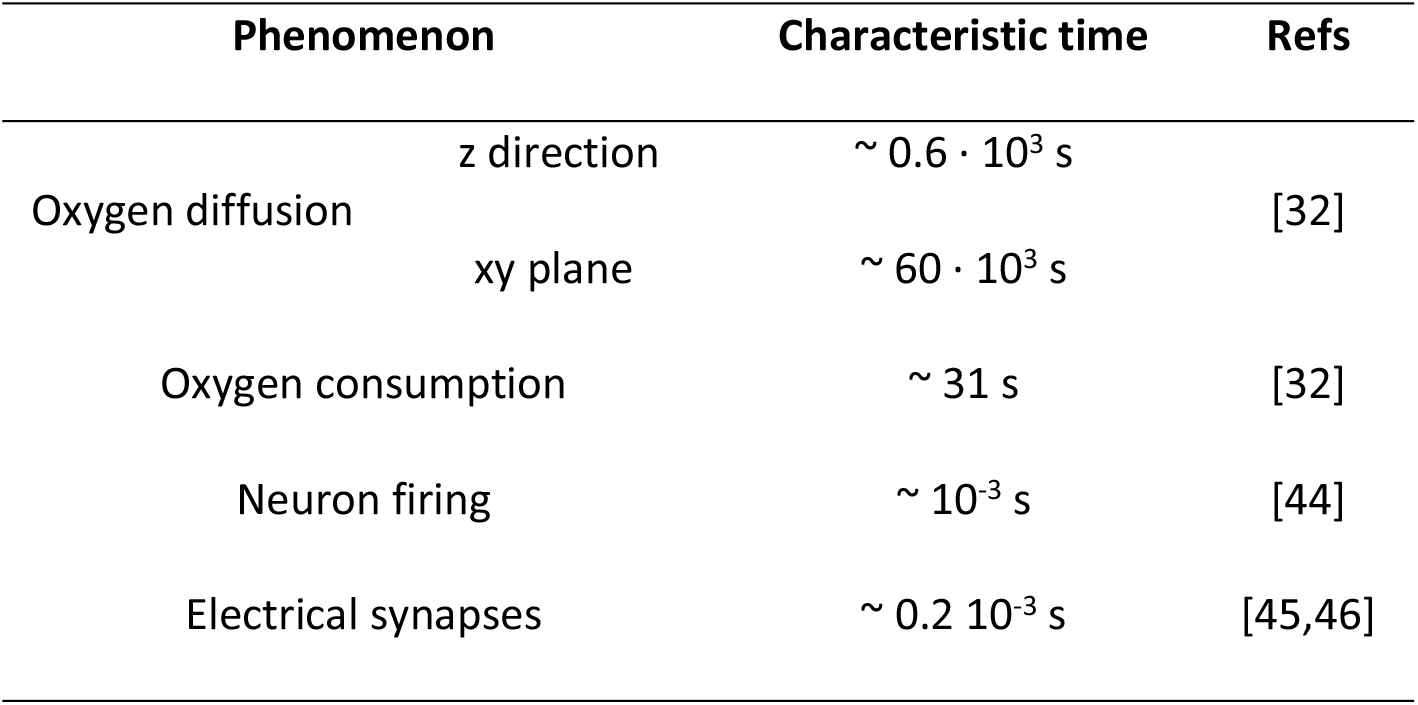
Characteristic times of the phenomena involved.

### 2.2 Model of oxygen and single-neuron firing

The model of oxygen and single-neuron firing describes the consumption of oxygen by the neuron for maintaining vital functions and for firing. Since O_2_ consumption generates an oxygen concentration gradient from the top (z=h) to the bottom (z=0) of the medium, the governing equation of a single neuron in a column of aqueous medium is given by:

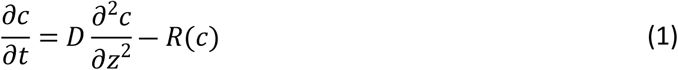

where *c* is the oxygen concentration, D is the diffusion constant of oxygen in water and *R*(*c*) is the oxygen consumption. Since 75% of oxygen is devoted to fuel neuronal spiking activity and the remaining 25% to sustain vital cell functions [27,28], *R*(*c*) can be written as:

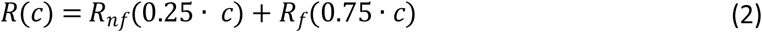

where *R*_*nf*_(*c*) is the rate at which oxygen is consumed for cellular and sub-cellular processes not directly linked to electrophysiological activity and *R*_*f*_(*c*) is the oxygen consumption contributing to neuron firing. *R*_*nf*_(*c*) can be described via Michaelis-Menten kinetics [32,50]:

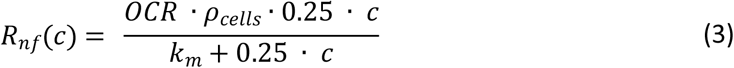

where *OCR* is the maximal oxygen consumption rate per cell, *ρ*_*cells*_ is the cell density and *k*_*m*_ is the Michaelis-Menten constant.

On the other hand, O_2_ consumption related to firing is as described by Wei et al., 2014 [37]:

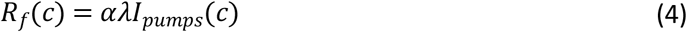

where *I*_*pumps*_ is the transport rate of ions across the membrane, measured in *mM*/*s, α* is the conversion factor from *mM*/*s* to *mg*/*l* and *λ* is the relative cell density. A detailed description of the model and a list of the constants – typical of mammalian cells - and the respective units of measure are reported in the S1 Text, S1 Fig. and S1 Table.

### 2.2 Connectivity model

The neuronal network can be generated connecting the neurons with the desired positioning and wiring. Although the communication between neural cells is perceived both via electrical and chemical signalling [51], in this study we implemented the simplest model of the neuron-to-neuron signal transmission, i.e., via electrical coupling. Given a post-synaptic neuron *i* and assuming that it receives inputs from *N* pre-synaptic neurons *j*, its membrane potential is described by the following equation:

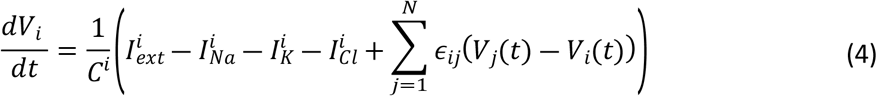

where *ϵ*_*ij*_ is the synaptic weight of the connection between the pre-synaptic neuron *j* and the post-synaptic neuron *i*, with the dimension of a conductance. In this work, we assumed that each electrical synapse has a weight inversely proportional to the distance between the two connected neurons [52]. It has been observed that the structure of neuronal networks of both brain areas and cellular monolayers can be described by Small-World (SW) graphs [40,43,53]. SW networks are topological structures with short characteristic path length *L* and high clustering coefficient *CC* resulting in a small-worldness *σ* greater than one [54,55]. Details and definitions of the SW metrics are provided in S2 Table.

To allow the generation of networks with any size and wiring patterns, the model of oxygen and single neuron firing is integrated in a customized Simulink library (Fig 2). The following parameters can be user-defined: i) the boundary oxygen concentration *c*_*0*_ on top of the medium, ii) the height of the medium *h*, and iii) the metabolic and firing parameters of the neuronal cell.

**Fig 2.**
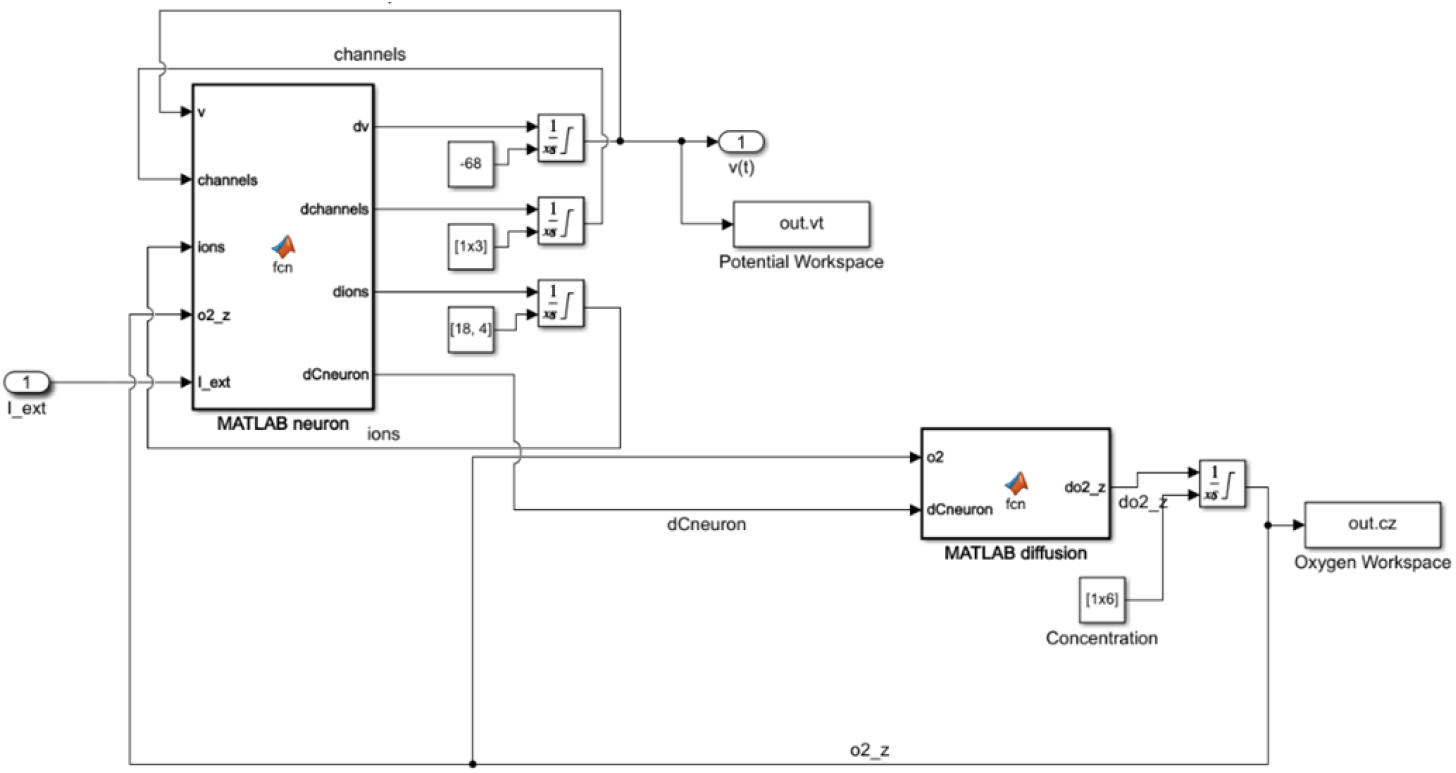
Simulink scheme of the single-neuron model. Simulink blocks embedding functions describing oxygen diffusion through the column medium (MATLAB diffusion) and neuron firing dynamics (MATLAB neuron).

Then, the user can express the wiring information through an adjacency matrix, usually used to describe the neuron-neuron connections [53,56,57]. Starting from the number of nodes and links and the metrics characterizing the networks, the adjacency matrix can be obtained using the Watts-Strogatz method [50]. Given the entries of the adjacency matrix, i.e., the weights *ϵ*_*ij*_, the *Digitoids* are generated automatically.

## 3. Materials and Methods

### 3.1 Single neuron simulations

For assessing the influence of oxygen availability on neuron firing, firstly the models of oxygen and single neuron firing described in Section 2.2 were computed, varying both the boundary oxygen concentrations *c*_0_ stepwise from 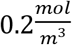 (i.e., the maximum available oxygen concentration in water) to 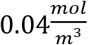 (i.e., the critical oxygen concentration for cell survival) [50], and the height of the medium above the cell at *h* = 0.5 *mm*, 1 *mm* and 3 *mm* (corresponding to the experimental volumes of medium usually used for neuron electrophysiological recordings [58–60]). All the parameter combinations were simulated for 10 seconds.

### 3.2 Output validation of Digitoids

The performance of the *Digitoids* was evaluated against experimental data, following the pipeline shown in Fig 3. Specifically, we compared the simulations generated by the platform with outcomes from the *in vitro* neuronal networks presented in Ballesteros-Esteban et al. [60]. The authors describe the morphology and electrophysiological activity of 19 *in vitro* neuronal networks derived from invertebrate neurons. They tracked the networks through imaging and MEA recordings for three weeks, establishing SW connectivity graphs. A detailed summary of the experimental procedure adopted by the authors is reported in S2 Text. Here, we exploited the SW metrics identified from day in vitro (DIV) 11 to DIV 16 to sketch the corresponding *Digitoids* and simulate their dynamics for the same duration as the recordings performed by Ballesteros-Esteban et al. [60].

**Fig 3.**
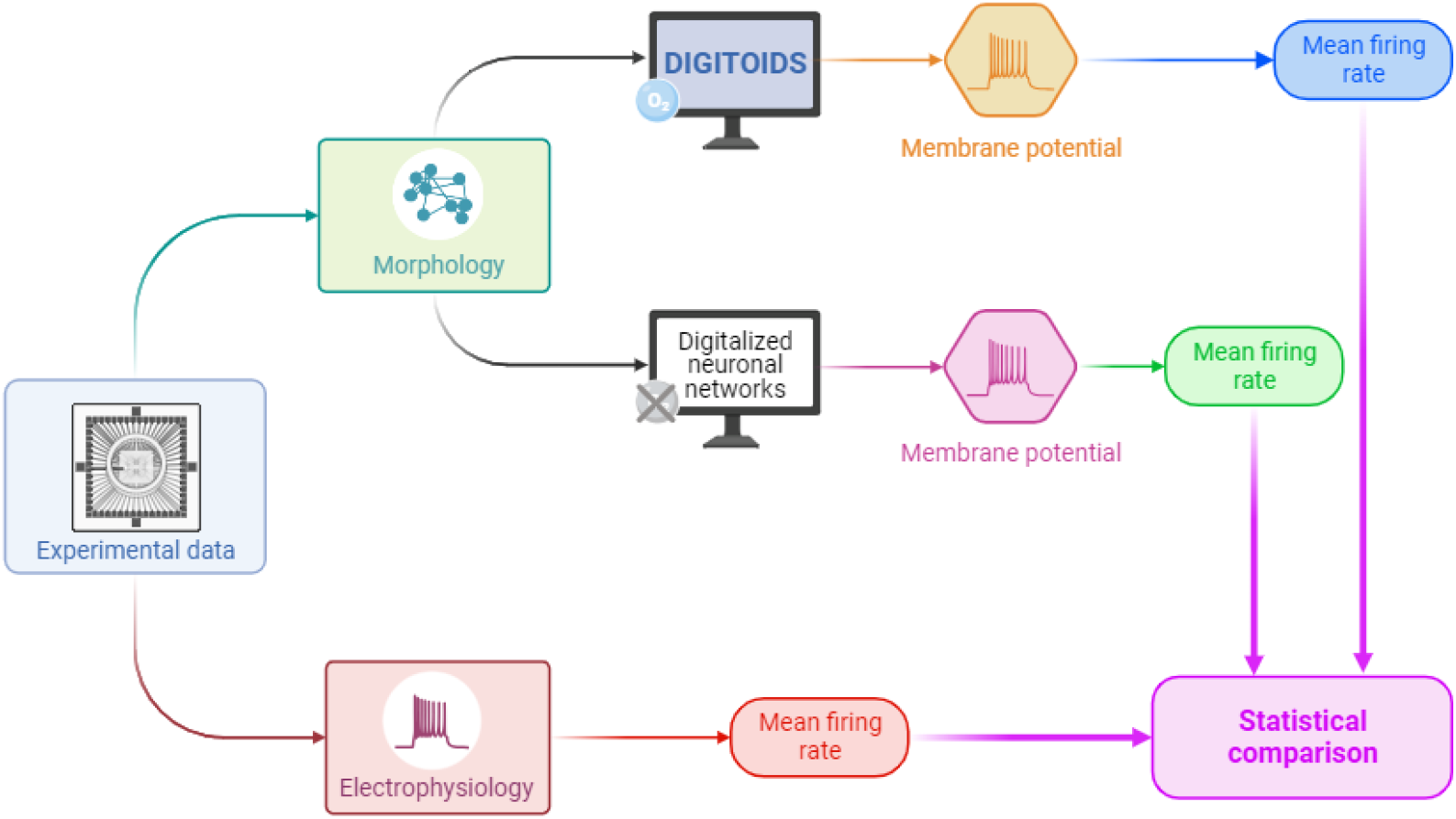
Validation pipeline. Schematic workflow of the validation pipeline adopted for testing the Digitoids’ performance with experimental data.

The output signals generated by the *Digitoids* were then processed for detecting the spikes with the same adaptive threshold as in Ballesteros-Esteban et al. [60]. The neuron firing rate was calculated by dividing the number of spikes for the total time of simulation. Finally, the mFR (mean Fring Rate) over the networks was extracted and compared with the values reported by Ballesteros-Esteban and co-workers. To demonstrate the importance of oxygen dependence in modelling neuron firing through equations(1)-(4), the same layouts were also simulated using the standard Hodgkin-Huxley model (equations S.2) which does not consider the relation occurring between the ion pump activity and the local oxygen concentration.

### 3.3 Statistical analysis

Statistical analyses were performed using GraphPad Prism 8 (GraphPad Software, Boston, Massachusetts USA). Since the hypothesis of a Gaussian distribution of the mFR from the experimental and *in silico* data was not confirmed by the Shapiro-Wilk test for normality, the non-parametric Kruskal-Wallis test was used for assessing the *Digitoids’* performance. Specifically, we determined the extent to which the oxygen-dependent firing model in equations (1)-(4) is better at replicating the *in vitro* counterpart than the conventional model. The significance level was set to 0.05 for both normality test and comparison.

## 4. Results

### 4.1 Dependence of firing on oxygen availability

Fig 4 shows a detail of the neuron membrane potential for different values of *c*_*0*_ (at z=h) and the rate of oxygen decrease at the neuron level (z=0) during firing. The graphs clearly show the dependence of firing on local oxygen concentration. The membrane voltage and the oxygen concentration at z=0 for all parameters assessed are reported in S2-S6 Fig.

**Fig 4.**
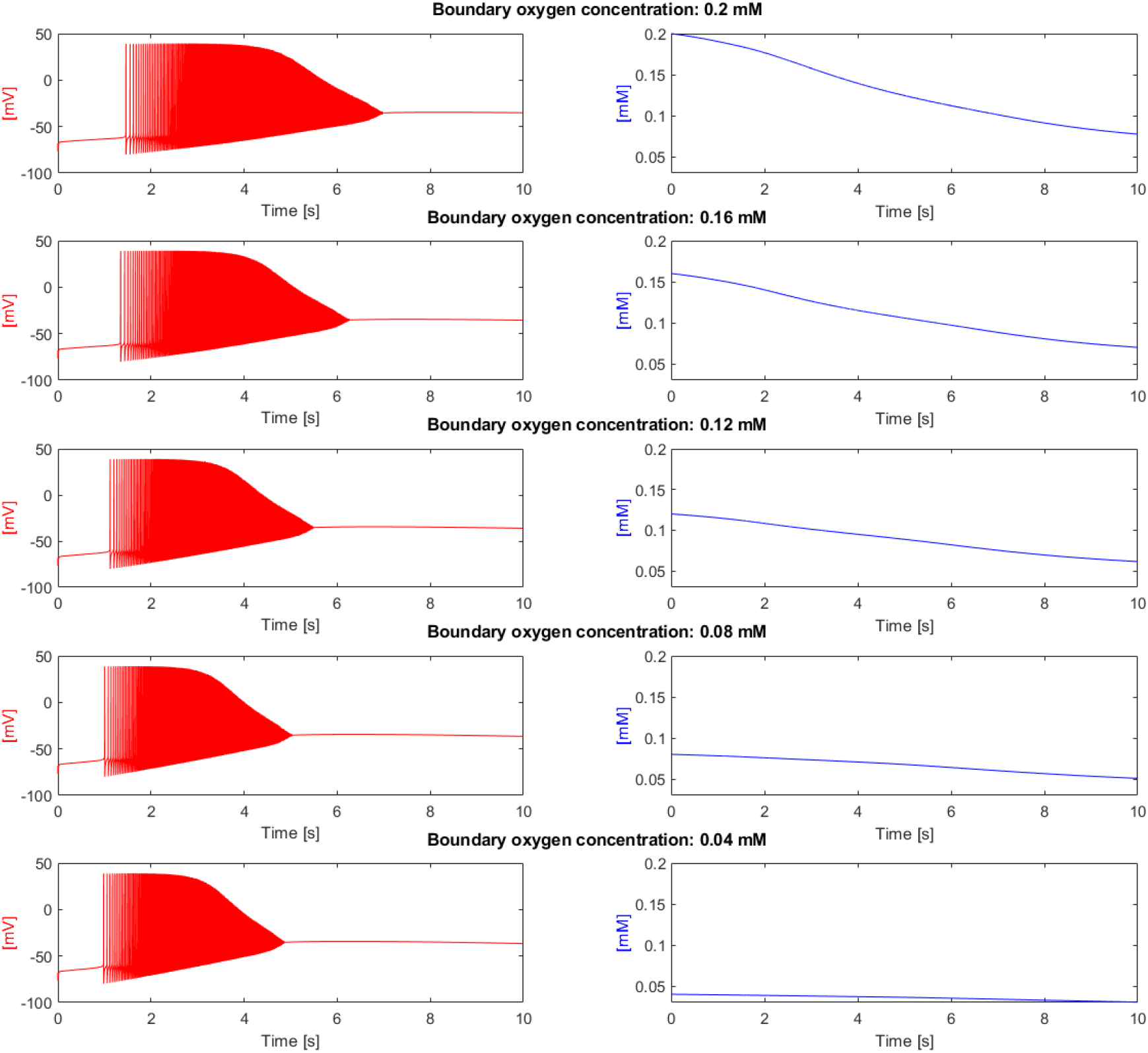
Single neuron output. Membrane potential (left column) expressed in mV and oxygen concentration at the neuron level (right column) expressed in mM, over 10 seconds with h=3 mm and different boundary oxygen concentrations c0.

Notably, the duration of a spike train (i.e., a sequence of action potentials originating from a single neuron [61]) was sensitive to the level of oxygen available to the neuron [44]. We calculated the starting train time as the time at which the signal overcomes the threshold of -60 mV [62,63], and the ending train time as the time at which the derivative of the signal turns close to 0. The duration of the spike train, *t*_*train*_, was expressed as the difference between ending and starting times, reported in Fig 5, for the different values of *c*_*0*_ and the different media heights, *h*. As expected, *t*_*train*_ grows with higher *c*_*0*_ and with smaller values of *h*. It should also be noted that the variations between different values of *h* decrease with decreasing *c*_*0*_ (light blue region of the surface of Fig 5). This implies that firing behaviour is strongly limited by diffusion; below a certain threshold of oxygen concentration, firing is suppressed since it cannot be sustained [64–66]. For further information see S3 Table.

**Fig 5.**
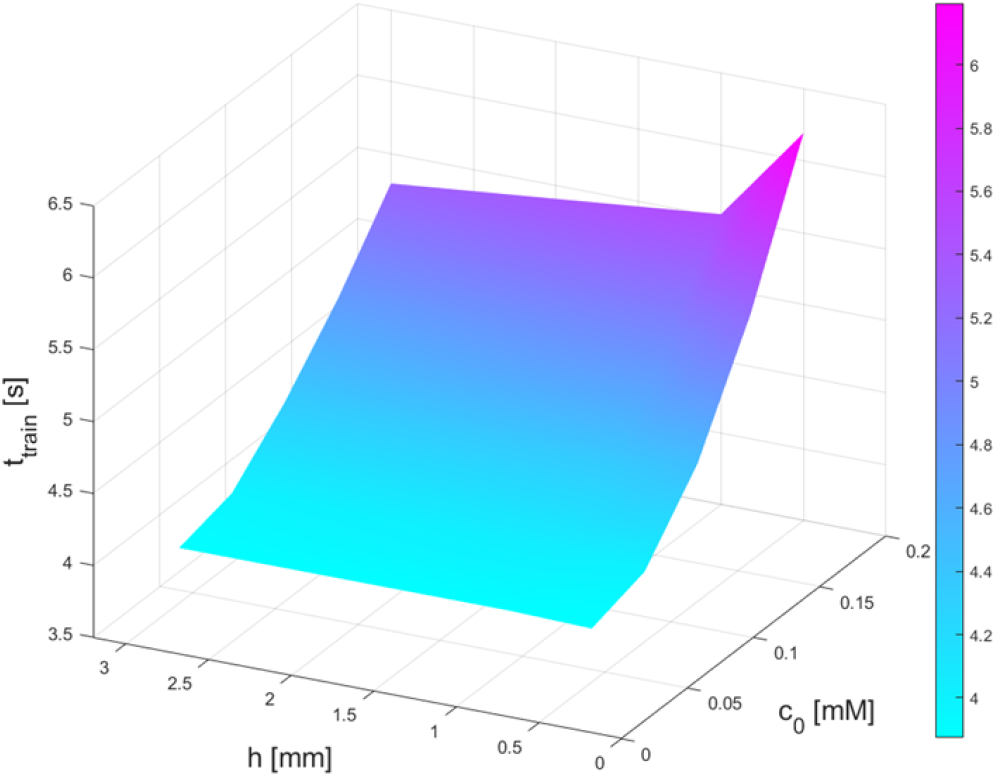
Duration of the first train of action potentials. The graph depicts the duration of the first train of action potentials varying the boundary oxygen concentration (c_0_) and the height of the medium height h.

It should also be noted that the rate of membrane depolarization increases as the available oxygen concentration at the neuron decreases. This behaviour was reported in both brain slices and *in vitro* cultures exposed to hypoxia [64,67–70], and can be explained by the fact that a reduced availability of oxygen causes a break in the homeostatic maintenance of ion concentrations between inner and outer neuron environments, which is responsible for sustaining the electrical activity of the neuron. Essentially, when oxygen levels are reduced, the Na^+^-K^+^-ATP pump lacks resources for fuelling ion transport. This situation leads to an accumulation of K^+^ ions outside the cell, i.e., [K^+^]_o_ increases (Fig 6), leading to membrane depolarization [66].

**Fig 6.**
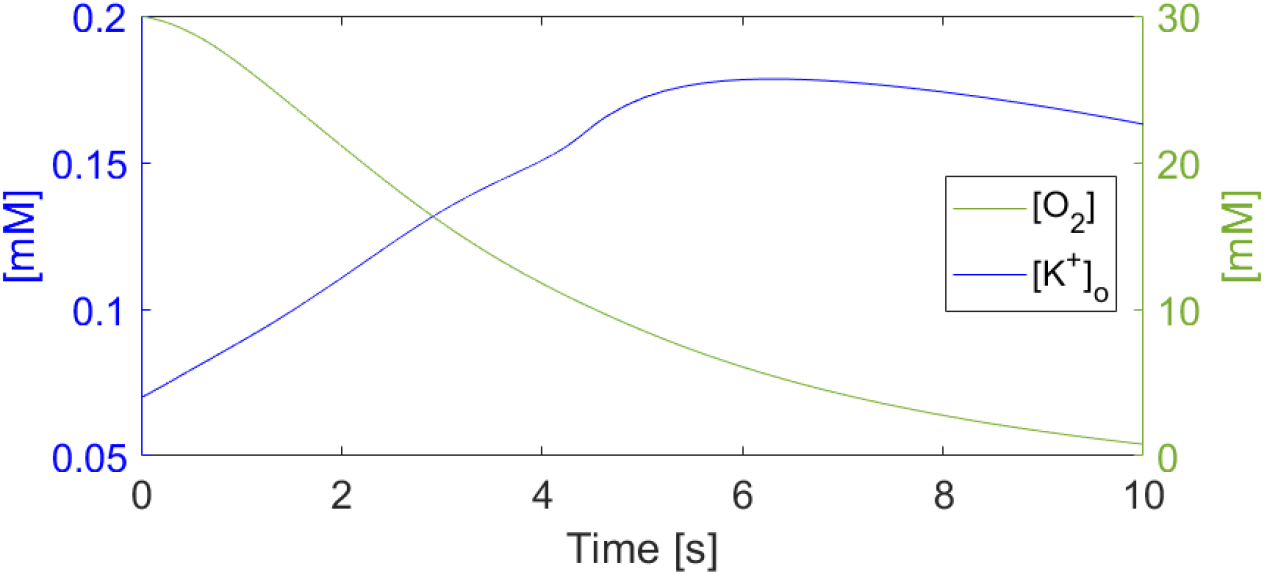
Potassium and oxygen concentrations. Trend over time of the extracellular K^+^ and O_2_ concentration

### 4.2 Digitoids’ performance

Fig 7 shows the mFR obtained from: i) the *in vitro* recordings reported in [60]; ii) the output from the corresponding *Digitoids*; and iii) simulations using the classic equations without the oxygen dependence on neuron firing activity. For all the DIV considered, no statistically significant differences were found between the mFR output from the *Digitoids* and the corresponding experimental data. On the other hand, the mFR output from the simulations of the classic oxygen-independent network models showed significant differences with the ones extracted *in vitro*.

**Fig 7.**
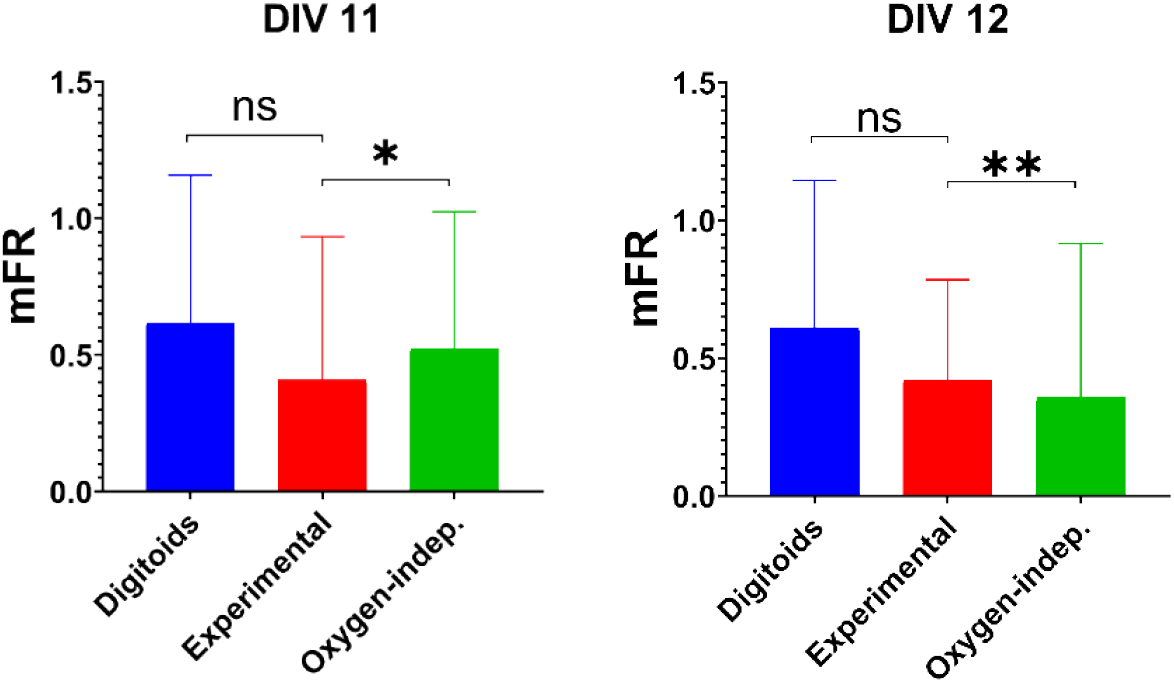
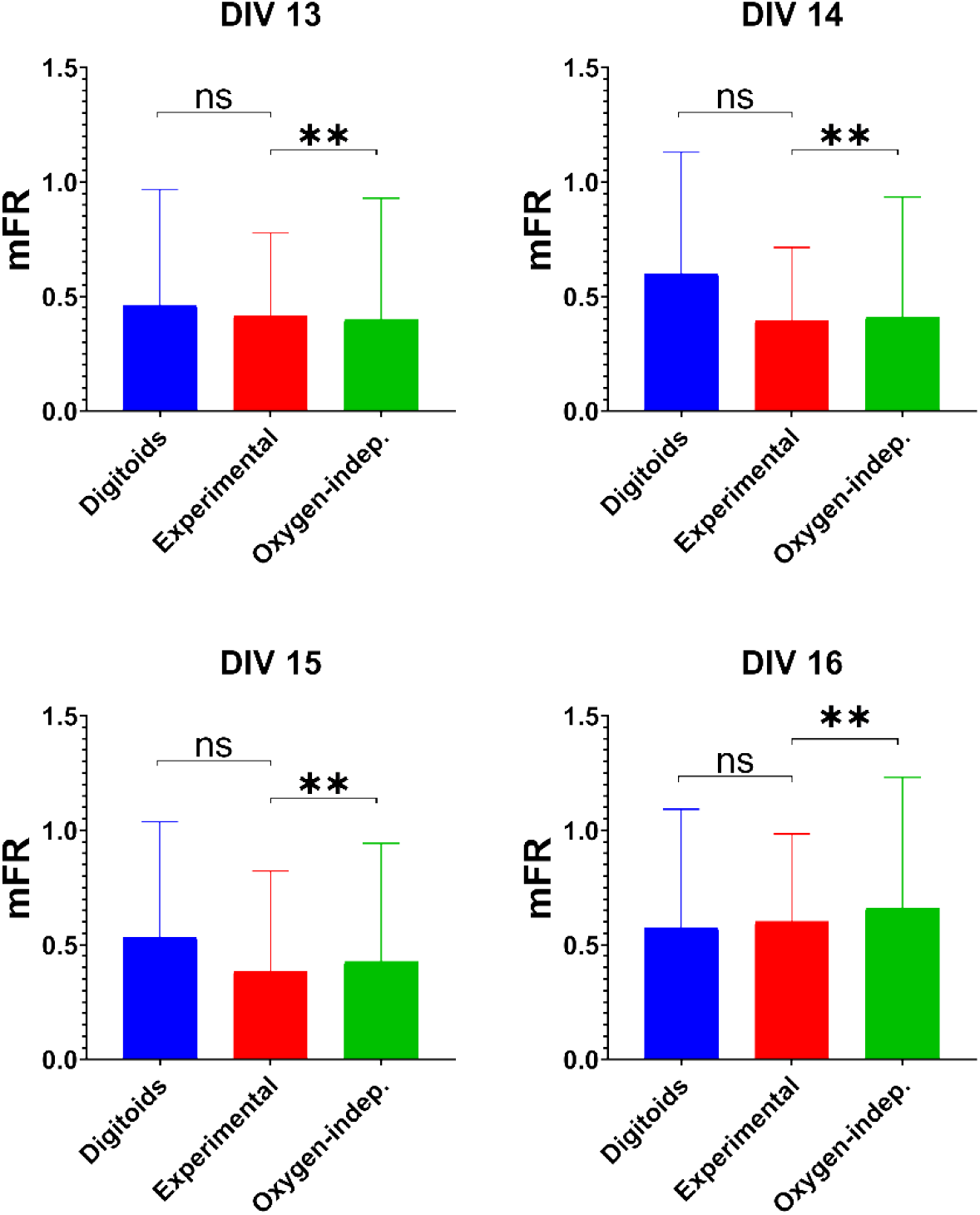
Validation results. Results of the statistical comparison of the mFR of the experimental data (in red), the Digitoids (in blue) and the oxygen-independent models (in green).

## 5. Discussion

There is plenty of evidence to show that oxygen levels are crucial to neuronal function *in vitro:* they significantly affect viability, oxidative stress and mitochondrial function [71]. However, the influence of O_2_ on *in vitro* electrophysiological behaviour is often neglected. In this work, we developed a computational platform which embeds a novel model of neuron firing where the oxygen dynamics of diffusion and consumption are introduced [71]. The platform can be used to design and simulate *Digitoids*, i.e., digital neuronal networks with the same layouts observed *in vitro*. The outputs are the membrane potentials and the oxygen concentration at the neuron level (z=0) over time. To validate the platform, we reproduced the metrics of *in vitro* neuronal networks seeded on commercial MEAs. No statistical differences were found between the mFR experimentally measured and those obtained with the *Digitoids*. To further assess the *Digitoids*’ performance, we also compared the *in vitro* mFRs with the classic Hodgkin-Huxley model applied to the same network layouts. The results are significantly different, demonstrating that local oxygen concentration and ion pump activity strongly affect electrophysiological dynamics.

It is worth highlighting that the experimental data derive from insect neurons, while the model parameters are typical of mammalian neurons. However, invertebrate neurons are widely used to advance knowledge about more complex organisms [72,73], since they exhibit morphological and electrophysiological features similar to mammalian neurons (e.g., mechanisms of membrane depolarization [70]).

A more in-depth validation of our platform would necessitate parallel recordings of electrophysiological and oxygen dynamics of *in vitro* neurons, along with a more exhaustive quantitative description of the network dynamics, rather than just its overall mean activity. Other metrics describing the physical and functional connections of neurons *in vitro* will be also taken into account [57].

The introduction of models of excitatory and inhibitory chemical synapses, and the integration of an analytical model of the oxygen demand for their function will be crucial to evaluate how the *Digitoids* behave in response to both electrical and chemical stimuli [74], as well as how neurons can reshape their network configuration to learn tasks, e.g., in a closed-loop approach [75,76]. The integration of model of synapses will enable a better understanding of signal integration and the ensuing dynamics, providing essential information for model construction and validation. Indeed, translation into models would require investigation of structural changes, receptors, and biochemical pathways. Our future efforts will focus on collecting more mechanistic data of the *in vitro* biological constructs to quantitively describe and integrate them within the *Digitoids*.

Thanks to its modularity, the platform delivers a flexible digital tool that can easily be adapted for mimicking *in vitro* models with increasing complexity (e.g., co-cultures, including other neural phenotypes). Since the user can easily modify the single neuron model as well as the culture settings and the network layouts, our approach can be customized for modelling neurospheres and cerebral organoids. Indeed, it is well known that insufficient nutrient supply, and the subsequent generation of non-viable cores, hinder the creation of mature traits and of a three-dimensional neural network [4].

While the concept of *digital twins* is progressively taking root in healthcare applications [77,78], their potential has yet to be explored for *in vitro* systems. For instance, digital twins could be used to improve the characterization, optimization and routine manufacturing of tissue culture technology and to replace animal testing, in compliance with the 3Rs (i.e., Reduction, Refinement and Replacement) principle [79]. The *Digitoids* represent the first step towards virtual copies of *in vitro* neuronal networks, as “digital twins” that could be used to support, or even replace, primary neuronal cultures, being easier to prepare and maintain, having longer lifespans and allowing high-throughput experiments [80].

## Supporting information captions

**S1 Text. Model of oxygen and single-neuron dynamics**. Detailed description of the mathematical model adopted for describing the oxygen diffusion/consumption and the single neuron dynamics.

**S1 Fig. Pump rate description**. Trend of membrane pump rate as a function of the oxygen input to the neuron for firing. c_f_=0.75·c is the oxygen percentage devoted to firing.

**S1 Table. Parameter inputs to the computational platform**.

**S2 Table. Small World metrics**. The network metrics exploited for describing the wiring pattern of a network composed of n nodes and m links. aij defines the connection state of nodes i and j: it is 1 if nodes i and j are connected and it is 0 vice versa.

**S2 Text. Experimental data description**. Detailed description of the experimental settings ad experiments conducted to produce the electrophysiological and morphological data compared against the *Digitoids*’s output.

**S3 Table. Duration of the first train of action potentials, varying medium height (first column) and boundary oxygen concentration (first row)**.

**S2-S4 Fig. Single neuron simulation with h= 3 mm, 1 mm and 0.5 mm**. Membrane potential profile: left side, red trace; oxygen concentration profile: right side, blue trace.

**S5-S6 Fig. Detail of the first 10 s of simulation with h= 1 mm and 0.5 mm**. Membrane potential profile: left side, red trace; oxygen concentration profile: right side, blue trace.

